# SARS-CoV-2 vaccination induces neutralizing antibodies against pandemic and pre-emergent SARS-related coronaviruses in monkeys

**DOI:** 10.1101/2021.02.17.431492

**Authors:** Kevin O. Saunders, Esther Lee, Robert Parks, David R. Martinez, Dapeng Li, Haiyan Chen, Robert J. Edwards, Sophie Gobeil, Maggie Barr, Katayoun Mansouri, S. Munir Alam, Laura L. Sutherland, Fangping Cai, Aja M. Sanzone, Madison Berry, Kartik Manne, Anyway B. Kapingidza, Mihai Azoitei, Longping V. Tse, Trevor D. Scobey, Rachel L. Spreng, R. Wes Rountree, C. Todd DeMarco, Thomas N. Denny, Christopher W. Woods, Elizabeth W. Petzold, Thomas H. Oguin, Gregory D. Sempowski, Matthew Gagne, Daniel C. Douek, Mark A. Tomai, Christopher B. Fox, Robert Seder, Kevin Wiehe, Drew Weissman, Norbert Pardi, Priyamvada Acharya, Hanne Andersen, Mark G. Lewis, Ian N. Moore, David C. Montefiori, Ralph S. Baric, Barton F. Haynes

## Abstract

Betacoronaviruses (betaCoVs) caused the severe acute respiratory syndrome (SARS) and Middle East Respiratory Syndrome (MERS) outbreaks, and now the SARS-CoV-2 pandemic. Vaccines that elicit protective immune responses against SARS-CoV-2 and betaCoVs circulating in animals have the potential to prevent future betaCoV pandemics. Here, we show that immunization of macaques with a multimeric SARS-CoV-2 receptor binding domain (RBD) nanoparticle adjuvanted with 3M-052-Alum elicited cross-neutralizing antibody responses against SARS-CoV-1, SARS-CoV-2, batCoVs and the UK B.1.1.7 SARS-CoV-2 mutant virus. Nanoparticle vaccination resulted in a SARS-CoV-2 reciprocal geometric mean neutralization titer of 47,216, and robust protection against SARS-CoV-2 in macaque upper and lower respiratory tracts. Importantly, nucleoside-modified mRNA encoding a stabilized transmembrane spike or monomeric RBD protein also induced SARS-CoV-1 and batCoV cross-neutralizing antibodies, albeit at lower titers. These results demonstrate current mRNA vaccines may provide some protection from future zoonotic betaCoV outbreaks, and provide a platform for further development of pan-betaCoV nanoparticle vaccines.

## MAIN

The emergence of three betacoronavirus (BetaCoV) outbreaks over the last 18 years necessitates the development of countermeasures that can prevent future pandemics ^1–4^. SARS-CoV-1 and SARS-CoV-2 (betaCoV group 2b) and MERS (betaCoV group 2c) emerged from cross-species transmission events where humans were infected with bat or camel CoVs ^5–9^. In particular, betaCoVs that are genetically similar to SARS-CoV-1 and SARS-CoV-2 and bind to the human ACE2 receptor for virus entry circulate in civets, bats, and Malayan pangolins ^5,6,10–12^. Thus, SARS-related animal coronaviruses represent betaCoVs that can be transmitted to humans. Neutralizing antibodies can prevent or treat betaCoV infection and represent potential countermeasures against current human betaCoVs and pre-emergent viruses ^13–22^. Cross-neutralizing antibodies capable of neutralizing multiple betaCoVs, have been isolated from SARS-CoV-1 infected humans ^13,23,24^, providing proof-of-concept for development of betaCoV vaccines against group 2b *Sarbecoviruses* ^25^. A critical target of cross-neutralizing antibodies is the receptor binding domain (RBD) ^22,23,25^. RBD immunogenicity can be augmented by arraying multiple copies of it on nanoparticles, mimicking multimeric virus-like particles ^26–30^. Vaccine induction of cross-neutralizing antibodies has been reported in *in vitro* neutralization assays against CoV pseudoviruses in mice ^29,30^. However, it is unknown whether spike vaccination of primates can elicit cross-neutralizing betaCoV antibodies against SARS-CoV-1, bat betaCoVs, or against SARS-CoV-2 escape viruses. Here, we demonstrate the ability of a SARS-CoV-2 RBD 24-mer subunit nanoparticle vaccine—produced with an easily modifiable sortase-ferritin platform^31^—to elicit in monkeys potent cross-neutralizing antibodies against SARS-CoV-1, SARS-CoV-2, SARS-CoV-2 variant B.1.1.7, as well as SARS-related bat betaCoVs. RBD nanoparticle vaccination protected monkeys against SARS-CoV-2 after respiratory challenge. Lastly, we show lipid nanoparticle-encapsulated Spike mRNA vaccines similar to those in clinical use elicit cross-neutralizing CoV antibodies, albeit at lower titers.

We and others have demonstrated the SARS-CoV-2 RBD has an epitope to which broadly cross-reactive neutralizing antibodies can bind ^13,24,32^, and the DH1047 RBD antibody cross-neutralizes SARS-CoV-1, CoV-2 and bat CoVs^24^.To focus the immune response on cross-reactive neutralizing epitopes on betaCoVs, we designed a 24-mer SARS-CoV-2 RBD-ferritin nanoparticle vaccine. The RBD nanoparticle was constructed by first expressing recombinant SARS-CoV-2 RBD with a C-terminal sortase A donor sequence. Next, we expressed the 24-subunit, self-assembling protein nanoparticle *Helicobacter pylori* ferritin with an N-terminal sortase A acceptor sequence ^31^. The RBD and *H. pylori* ferritin nanoparticle were conjugated together by a sortase A reaction (**Fig. 1a, Extended data Fig. 1**) ^31^. Analytical size exclusion chromatography and negative stain electron microscopy confirmed that RBD was conjugated to the surface of the ferritin nanoparticle (**Fig. 1a**, **Extended data Fig. 1b,c).** The RBD sortase A conjugated nanoparticle (RBD-scNP) bound to human ACE2, the receptor for SARS-CoV-2, and to potently neutralizing RBD antibodies DH1041, DH1042, DH1043, DH1044, and DH1045 ^24^ (**Fig. 1b**). The epitopes of these antibodies are focused on the receptor binding motif within the RBD ^24^. Cross-neutralizing antibody DH1047 also bound to the RBD-scNP (**Fig. 1b**). The RBD scNP lacked binding to SARS-CoV-2 spike antibodies that bound outside of the RBD (**Fig. 1b**).

**Figure 1.**
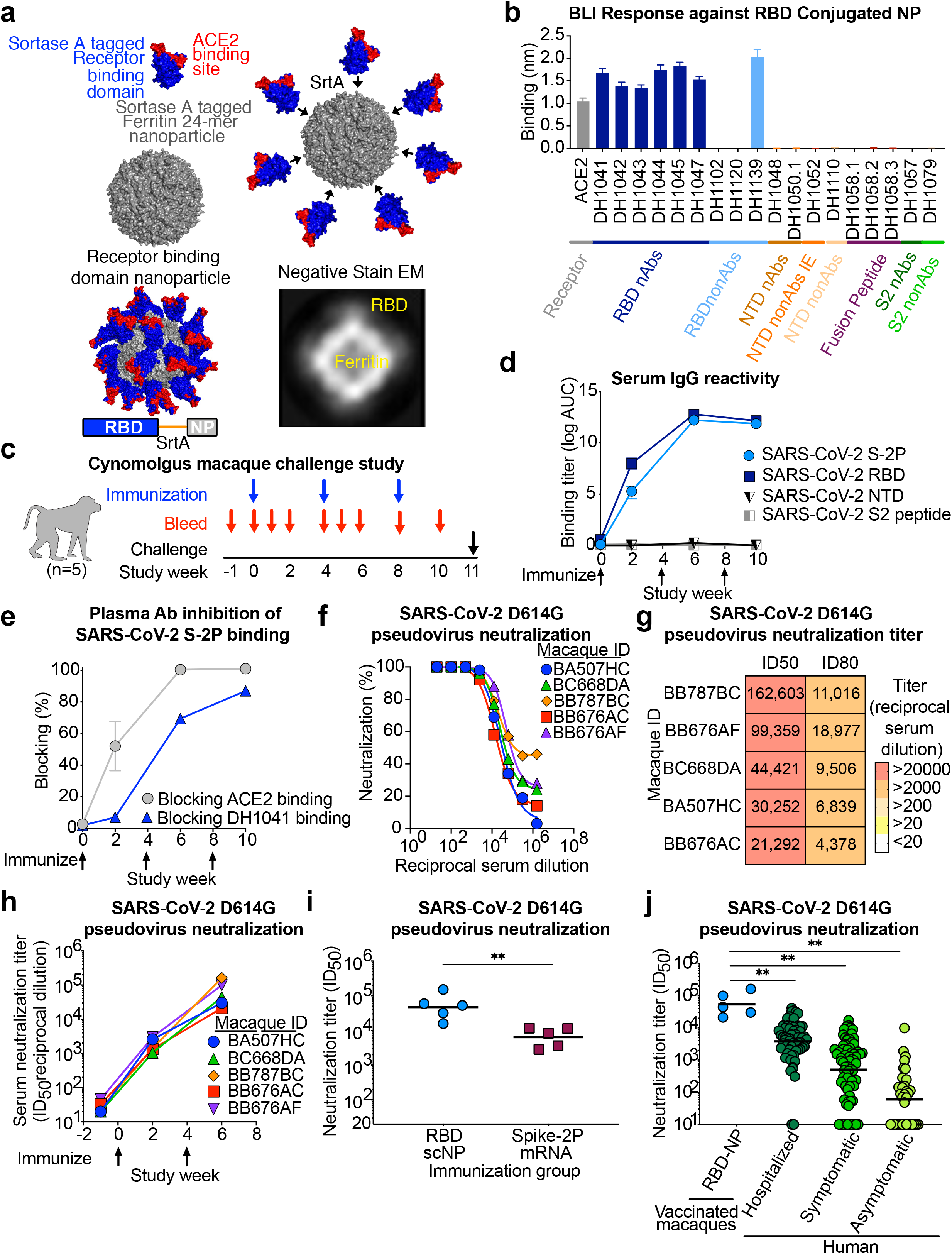
SARS-CoV-2 receptor binding domain (RBD) sortase conjugated nanoparticles (scNPs) elicits extremely high titers of SARS-CoV-2 pseudovirus neutralizing antibodies. **a.** SARS-CoV-2 RBD nanoparticles were constructed by expressing RBD with a C-terminal sortase A donor sequence (blue and red) and a *Helicobacter pylori* ferritin nanoparticle with N-terminal sortase A acceptor sequences (gray) on each subunit (top left). The RBD is shown in blue with the ACE2 binding site in red. The RBD was conjugated to nanoparticles by a sortase A (SrtA) enzyme conjugation reaction (top right). The resultant nanoparticle is modeled on the bottom left. Nine amino acid sortase linker is shown in orange. Two dimensional class averages of negative stain electron microscopy images of actual RBD nanoparticles are shown on the bottom right. **b.** Antigenicity of RBD nanoparticles determined by biolayer interferometry against a panel of SARS-CoV-2 antibodies and the ACE2 receptor. Antibodies are color-coded based on epitope and function. N-terminal domain (NTD), nonAbs IE, infection enhancing non-neutralizing antibody; nAb, neutralizing antibody; nonAb, non-neutralizing antibody. Mean and standard error from 3 independent experiments are shown. **c.** Cynomolgus macaque challenge study scheme. Blue arrows indicate RBD-NP immunization timepoints. Intranasal/intratracheal SARS-CoV-2 challenge is indicated at week 10. **d.** Macaque serum IgG binding determined by ELISA to recombinant SARS-CoV-2 stabilized Spike ectodomain (S-2P), RBD, NTD, and Fusion peptide (FP). Binding titer is shown as area-under-the curve of the log_10_-transformed curve. Arrows indicate immunization timepoints. **e.** Plasma antibody blocking of SARS-CoV-2 S-2P binding to ACE2-Fc and RBD neutralizing antibody DH1041. Group mean and standard error are shown. **f.** Dose-dependent serum neutralization of SARS-COV-2 pseudotyped virus infection of ACE2-expressing 293T cells. Serum was collected after two immunizations. The SARS-CoV-2 pseudovirus spike has an aspartic acid to glycine change at position 614 (D614G). Each curve represents a single macaque. **g.** Heat map of serum neutralization ID50 and ID80 titers for SARS-COV-2 D614G pseudotyped virus after two immunizations. **h.** SARS-COV-2 D614G pseudotyped virus serum neutralization kinetics. Each curve represents a single macaque. **i.** Comparison of serum neutralization ID50 titers from cynomolgus macaques immunized with recombinant protein RBD nanoparticles (blue) or nucleoside-modified mRNA-LNP expressing S-2P (burgundy) (\*\**P*<0.01, Two-tailed Exact Wilcoxon test n = 5). **j.** Comparison of serum neutralization titers obtained from RBD-scNP-vaccinated macaques (blue) and SARS-CoV-2 infected humans (shades of green). Human samples were stratified based on disease severity as asymptomatic (N=34), symptomatic (n=71), and hospitalized (N=60) (\*\**P*<0.01, Two-tailed Wilcoxon test n = 5).

To assess immunogenicity of the RBD scNP, we immunized five cynomolgus macaques three times intramuscularly four weeks apart (**Fig. 1c**). Immunogenicity of the protein was adjuvanted with the TLR7/8 agonist 3M-052 absorbed to alum ^33^. After a single immunization, all five macaques generated binding IgG antibodies against SARS-CoV-2 RBD and stabilized Spike ectodomain (S-2P) (**Fig. 1d**). Boosting once maximally increased plasma SARS-CoV-2 RBD and S-2P-specific IgG antibody titers (**Fig. 1d**). ACE2 competitive binding assays demonstrated the presence of antibodies against the ACE2 binding site within the receptor binding domain. Plasma antibodies blocked the ACE2 binding site on SARS-CoV-2 S-2P by 52% after one immunization and 100% after two immunizations (**Fig. 1e**). Similarly, plasma antibodies blocked the binding of ACE2-binding site-focused, RBD neutralizing antibody DH1041, although to a lesser degree than ACE2 (**Fig. 1e**). Vaccine induction of neutralizing antibodies was assessed against a SARS-CoV-2 pseudovirus with an aspartic acid to glycine substitution at position 614 (D614G)^34^. Two RBD scNP immunizations induced potent serum neutralizing antibodies (**Fig. 1f-h**). The fifty percent inhibitory reciprocal serum dilution (ID50) neutralization titers ranged from 21,292 to 162,603 (**Fig. 1g**). In cynomolgus macaques, serum neutralization titers against SARS-CoV-2 D614G pseudovirus elicited by RBD-scNP immunization were significantly higher than titers elicited by stabilized transmembrane (TM) Spike (S-2P) lipid-encapsulated nucleoside-modified mRNA (mRNA-LNP) immunization that is analogous to the immunogen and vaccine platform in the Moderna and Pfizer/BioNTech COVID-19 vaccines (**Fig. 1i,***P*<0.01 Exact Wilcoxon test, n=5)^35,36^. The group geometric mean ID50, measured as reciprocal serum dilution, for the macaques immunized with RBD-ScNP was 47,216 compared to 6,469 for mRNA-LNP immunized macaques. When compared to natural infection, RBD-scNP vaccination elicited ID50 neutralization titers higher than those elicited in humans with SARS-CoV-2 symptomatic infection, asymptomatic infection, or infection requiring hospitalization (**Fig. 1j**). Thus, RBD-scNP vaccination elicits significantly higher neutralizing titers compared to current vaccine platforms and natural infection.

The new SARS-CoV-2 variant B.1.1.7 is widespread in the United Kingdom (UK) and is spreading globally^37,38^. Moreover, the B.1.1.7 variant has been suggested to have higher infectivity and has mutations in the receptor binding domain that may limit neutralization efficacy of RBD-specific antibodies ^37,38^. Therefore, we tested the serum from the RBD-scNP-vaccinated macaques against a pseudovirus with B.1.1.7 spike. The macaque serum potently neutralized a pseudovirus bearing the Spike from the B.1.1.7 variant of SARS-CoV-2 (**Fig. 2a,b**). Similarly, neutralizing antibodies elicited by mRNA-LNP encoding the stabilized TM spike neutralized the B.1.1.7 variant of SARS-CoV-2 equally as well the D614G variant of SARS-CoV-2, albeit at titers below those observed with RBD-scNP immunization (**Fig. 2a,b**). Thus, the potentially more transmissible B.1.1.7 variant of SARS-CoV-2 was equally susceptible to vaccine-induced neutralizing antibodies as the SARS-CoV-2 D614G variant. Additionally, we determined RBD-scNP and S-2P mRNA-LNP immune plasma IgG binding to S ectodomain spike proteins containing mutations observed in strains circulating in Danish minks, as well as B1.351, P.1, and B.1.1.7 SARS-CoV-2 strains ^38–40^. As a control, we tested ACE2 receptor and SARS-CoV-2 cross-neutralizing antibody DH1047 binding to each spike mutant. The presence of D614G increased binding to spike by ACE2 and RBD cross-reactive antibody DH1047 (**Fig. 2c,d**). Similarly, plasma IgG from S-2P mRNA-LNP-immunized macaques and RBD-scNP-immunized macaques exhibited increased binding to spike in the presence of D614G (**Fig. 2c,d**). The addition of mutations from mink strains of SARS-CoV-2, the South African B1.351, Brazilian P.1 strain, and UK B.1.1.7 strains did not change binding magnitude for DH1047 nor macaque plasma IgG (**Fig. 2c,d**). However, the addition of these mutations slightly increased ACE2 binding compared to D614G alone (**Fig. 2c**). Next we tested binding to recombinant RBD monomers with and without the K417N, E484K, and N501Y mutations present in the B1.351 variant. ACE2 binding was decreased by K417N and increased by N501Y, while the RBD with all three mutations showed no change in ACE2 binding (**Fig. 2e**). RBD neutralizing antibody DH1041 is focused on the ACE2 binding site, and was knocked out by the E484K change (**Fig. 2e**). Importantly, the binding to SARS-CoV-2 RBD for cross-reactive antibody DH1047 was unchanged to RBDs with E484K or other mutations (**Fig. 2f**). Also, RBD-scNP-vaccinated macaque plasma IgG and S-2P mRNA-LNP-immunized macaque plasma IgG were unaffected by the RBD mutations (**Fig. 2e,f**). ^41^ Thus, RBD-scNP and mRNA-LNP-induced RBD binding antibodies were not sensitive to spike mutations present in neutralization-resistant UK, South Africa or Brazil SARS-CoV-2 variants.

**Figure 2.**
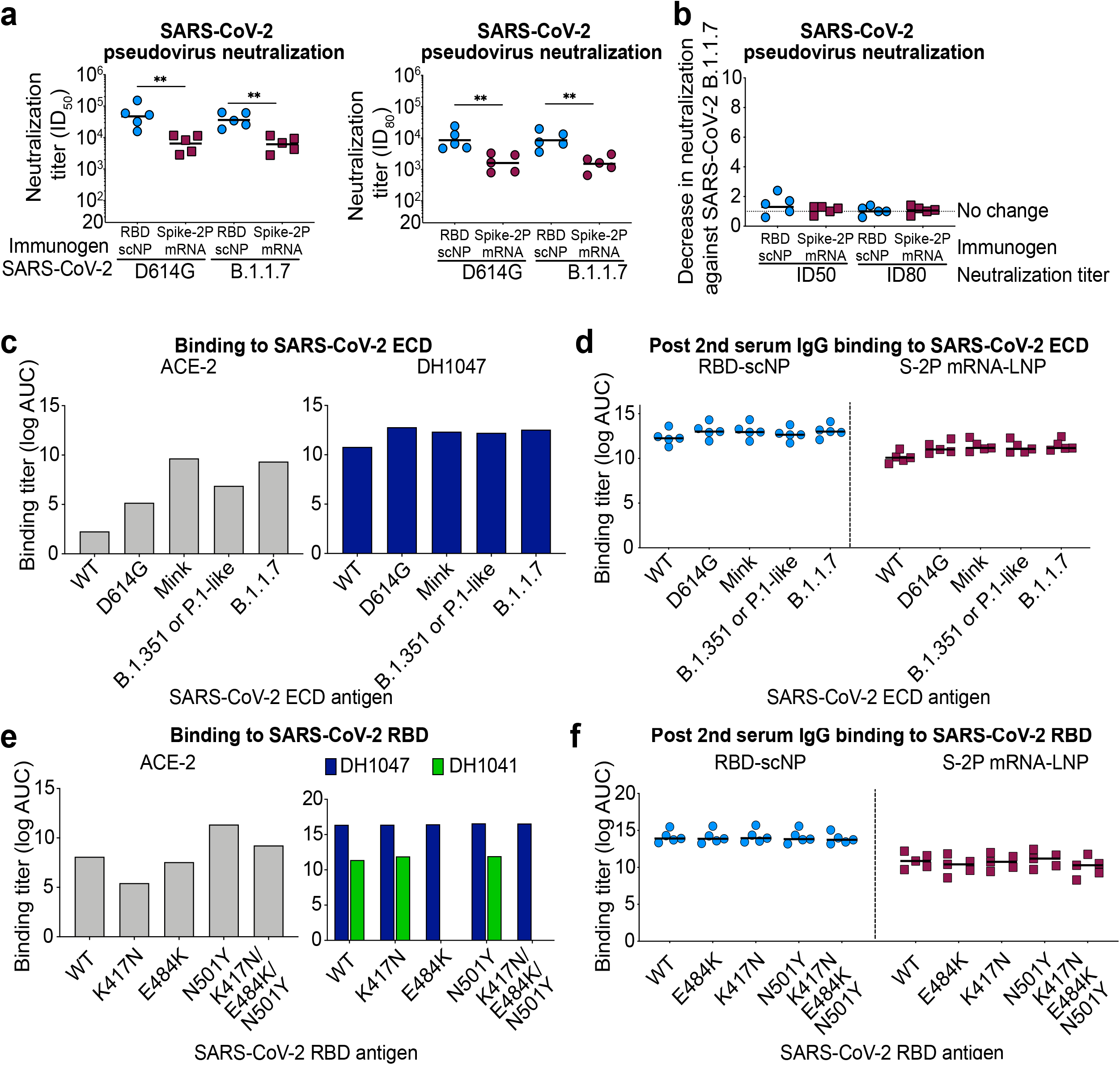
RBD-scNP immunization elicits higher titers of neutralizing antibodies and binding antibodies against more transmissible SARS-CoV-2 variants than stabilized spike mRNA-LNP vaccination. **a.** Comparison of serum neutralization ID50 titers from cynomolgus macaques immunized with recombinant protein RBD sortase conjugated nanoparticles (RBD-scNP; blue) or nucleoside-modified mRNA-LNP expressing S-2P (burgundy). ID50 titers (left) and ID80 titers (right) are shown. Horizontal bars are the group geometric mean (\*\**P*<0.01, Two-tailed Exact Wilcoxon test n = 5). **b.** Fold decrease in neutralization potency between neutralization of SARS-CoV-2 D614G and SARS-CoV-2 B.1.1.7 pseudoviruses. Fold change is shown for RBD-scNP-immunized and mRNA-LNP-immunized macaques based ID50 (left) and ID80 (right) titers. Horizontal bars are the group mean. **c.** ACE2 receptor and cross-neutralizing antibody DH1047 ELISA binding to SARS-CoV-2 Spike ectodomain (ECD) based on a Danish mink (H69/V70deI/Y453F/D614G/I692V), B.1.351 or P.1-like (K417N/E484K/N501Y/D614G), and B.1.1.7 (H69/V70del/Y144del/N501Y/A570D/D614G/P681H/T716I/S982A/D1118H) strains. Titers are shown as area under the log-transformed curve (log AUC). **d** RBD-scNP and S-2P mRNA-LNP-immunized macaque serum IgG ELISA binding to SARS-CoV-2 Spike variants shown in **c**. Serum was tested after two immunizations. Horizontal bars are the group mean. **e** ACE2 receptor (gray), cross-neutralizing antibody DH1047 (navy), and ACE2 binding site-targeting neutralizing antibody DH1041 (green) ELISA binding to SARS-CoV-2 Spike RBD monomers. RBD variants contain mutations found in circulating B.1.351 and P.1 virus strains. Titers are shown as area under the log-transformed curve (log AUC). **f** RBD-scNP and S-2P mRNA-LNP-immunized macaque serum IgG ELISA binding to SARS-CoV-2 Spike RBD variants shown in **e**. Serum was tested after two immunizations. Horizontal bars are the group mean.

While SARS-CoV-2 wild-type and variant neutralization remain a priority for halting the current pandemic, additional SARS-related CoVs that circulate in humans and animals remain a threat for future outbreaks ^42–44^. Therefore, we examined neutralization of SARS-CoV-1, SARS-related batCoV-WIV-1 and SARS-related batCoV-SHC014 by immune serum from vaccinated macaques^6,10,43,44^. After two immunizations, RBD-scNP, S-2P mRNA-LNP, and the RBD monomer mRNA-LNP elicited neutralizing antibodies against SARS-CoV-1, batCoV-WIV-1, and batCoV-SHC014 (**Fig. 3a, Extended data Fig. 2**). Neutralization was more potent for replication-competent SARS-CoV-2 virus compared to the other three SARS-related viruses (**Fig. 3a, Extended data Fig. 2**). Among the three immunogens, RBD-scNP elicited the highest neutralization titers and mRNA-LNP expressing monomer RBD elicited the lowest neutralization titers (**Fig. 3a, Extended data Fig. 2**). Also, RBD-scNP immunization elicited cross-reactive IgG binding against SARS-CoV-2, SARS-CoV-1, batCoV-RaTG13, batCoV-SHC014, pangolin CoV-GXP4L Spike proteins (**Fig. 3c** and **Extended data Figs. 3a,c**). RBD-scNP immune plasma IgG did not bind the S ectodomain of four endemic human CoVs, nor did it bind MERS-CoV S ectodomain (**Extended data figs. 3a,c)**. The lack of binding by plasma IgG to these latter five S ectodomains was consistent with RBD sequence divergence among groups 1, 2a, 2b and 2c coronaviruses (**Fig. 3e and Extended data figs. 4-6**). Nonetheless, the SARS-CoV-2 spike induced cross-reactive antibodies against multiple group 2b SARS-related betaCoVs, with the highest titers induced by RBD-scNP.

**Figure 3.**
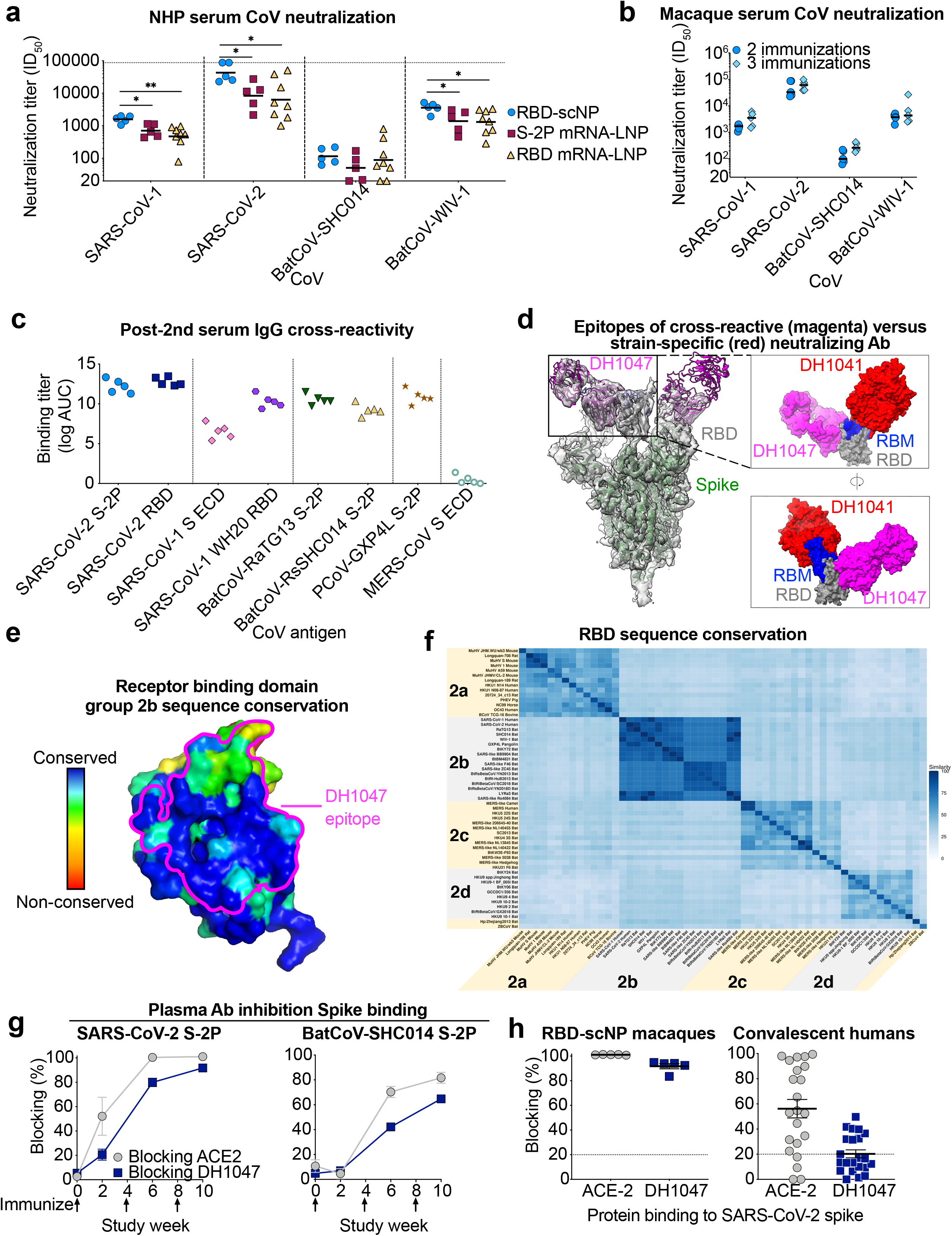
RBD-scNP vaccine induction of serum cross-neutralization of SARS-related betacoronavirus infection. **a.** Sera from macaques immunized twice with RBD-scNP, S-2P mRNA-LNP, or RBD mRNA-LNP neutralize replication-competent SARS-related human (SARS-CoV-1 and SARS-CoV-2) and bat (WIV-1 and SHC014) betaCoVs. Each symbol indicates the ID50 of an individual macaque. Horizontal bars are the group mean (\**P*<0.05, \*\**P*<0.01, Two-tailed Exact Wilcoxon test, n = 5 or 8). **b.** Comparison of serum cross-neutralization after two (circles) or three (diamonds) RBD scNP immunizations. Each symbol indicates the ID50 of an individual macaque. Horizontal bars are the group mean. **c.** Plasma IgG from macaques immunized twice with RBD-scNP binds to human, bat, and pangolin SARS-related betacoronavirus S protein in ELISA. ECD, ectodomain. **d.** Cryo-electron microscopy model comparing the epitopes of SARS-CoV-2-specific neutralizing RBD antibody (DH1041) and cross-neutralizing RBD antibody (DH1047). (Left) Cartoon view of Spike bound to DH1047 (magenta; PDB ID: 7LD1) fitted in the cryo-EM map (transparent gray; EMD-23279). Receptor Binding Domain (RBD) is in gray, Receptor Binding Motif (RBM) is in blue, and the rest of Spike is in forest green. (Top, right) Zoomed-in surface view of RBD (gray) and RBM (blue) binding interface with DH1047 (magenta) compared to the neutralizing RBD antibody DH1041 (red; PDB ID: 7LAA; EMD-23246). The RBDs of the two complexes from their respective cryo-EM structures were overlaid for comparison. (Bottom, right) A 180° rotated view of the top-right panel. **e.** DH1047 epitope is conserved within group 2b betaCoVs. Receptor binding domain (surface representation) colored by conservation within group 2b betacoronaviruses. DH1047 epitope is shown in magenta outline. **f.** Sequence similarity of RBD for representative betacoronaviruses. Heatmaps displaying pairwise amino acid sequence similarity for 57 representative betacoronaviruses. Dark blue shading indicates high sequence similarity. **g.** Macaque plasma antibody blocking of SARS-CoV-2 and batCoV-SHC014 S-2P binding to ACE2-Fc and RBD cross-neutralizing antibody DH1047. Group mean and standard error are shown. **h.** Rhesus plasma and convalescent human serum antibody blocking of SARS-CoV-2 S-2P binding to ACE2-Fc and RBD cross-neutralizing antibody DH1047. Each symbol represents a macaque or infected individual. Black horizontal bars indicate group mean and standard error.

The immune sera from RBD-scNP-immunized exhibited a similar cross-neutralizing profile as the cross-neutralizing antibody DH1047. DH1047 binds with <0.02 nM affinity to monomeric SARS-CoV-2 RBD (**Extended data fig. 3b**), and bound the RBD-scNP (**Fig. 1b**). DH1047 protects against SARS-CoV-2 infection in macaques and batCoV-WIV-1 infection in mice^24^; thus the DH1047 epitope is a principal betacoronavirus (CoV) cross-neutralizing antibody target for vaccination. The cross-reactive DH1047 epitope is adjacent to the N-terminus of the receptor binding motif distinguishing it from dominant RBM-focused neutralizing antibodies such as DH1041^24^ (**Fig. 3d**). Antibodies targeting near the DH1047 epitope would be predicted to be cross-reactive with group 2b betaCoVs given the high sequence conservation present in and immediately proximal to the DH1047 epitope (**Fig. 3e**). Comparison of RBD sequences showed them to be relatively conserved within betaCoV group 2b, but minimally conserved between groups 2b and 2c (**Fig. 3f and Extended data figs. 4-6**). To examine whether RBD-scNP-induced antibodies bound near the DH1047 epitope, we assessed plasma antibody blocking of DH1047 binding to SARS-CoV-2 S-2P ectodomain. Plasma from all RBD-scNP immunized macaques blocked the binding of ACE2 and DH1047 to SARS-CoV-2 S-2P ectodomain (**Fig. 3g and Extended data fig. 3d**). Since DH1047-blocking plasma antibodies could be SARS-CoV-2-specific, we also examined plasma IgG blocking of DH1047 binding to batCoV-SHC014. Similar to SARS-CoV-2 S binding, RBD-scNP plasma also blocked DH1047 binding to batCoV-SHC014 (**Fig. 3g**). While 96% of COVID-19 patients made antibodies that blocked ACE2 as a dominant RBD response, only 31% made antibodies that blocked the cross-reactive antibody, DH1047 (**Fig. 3h**). Thus, in natural SARS-CoV-2 infection the cross-reactive DH1047 IgG blocking response is subdominant, and RBD-scNP vaccination focused antibody responses to this subdominant cross-reactive neutralizing epitope.

Finally, to determine whether vaccination elicited coronavirus protective immunity, we challenged RBD scNP-vaccinated monkeys with 10^5^ plaque forming units of SARS-CoV-2 virus (**Fig 4a**). To assess virus replication in the upper and lower respiratory tract, E or N sgRNA was quantified in fluid from nasal swabs and bronchoalveolar lavage (BAL) two and four days after challenge (**Fig 4a**). In unimmunized macaques two days after challenge, there were on average 1.3×10^5^ and 1.2×10^4^ copies/mL of E gene sgRNA in the nasal swab and BAL fluids, respectively (**Fig. 4b,c**). In contrast, vaccinated monkeys had undetectable levels of subgenomic envelope E gene RNA in the upper and lower respiratory tract (**Fig. 4b,c**). We sampled monkeys again 2 days later to determine if detectable virus replication was present, but again found no detectable E gene sgRNA in any monkey BAL or nasal swab samples (**Fig. 4b,c**). Similarly, all RBD-scNP-vaccinated macaques had undetectable N gene sgRNA in BAL and the nasal swab fluid, except one macaque that had 234 copies/mL of N gene sgRNA detected on day 2 in nasal swab fluid (**Fig. 4d,e**). Viral replication was undetectable in this macaque by the fourth day after challenge (**Fig. 4d,e**). Thus, RBD-scNP-induced immunity prevented virus replication, and likely provided sterilizing immunity, in the upper and lower respiratory tract in all but one macaque.

**Figure 4.**
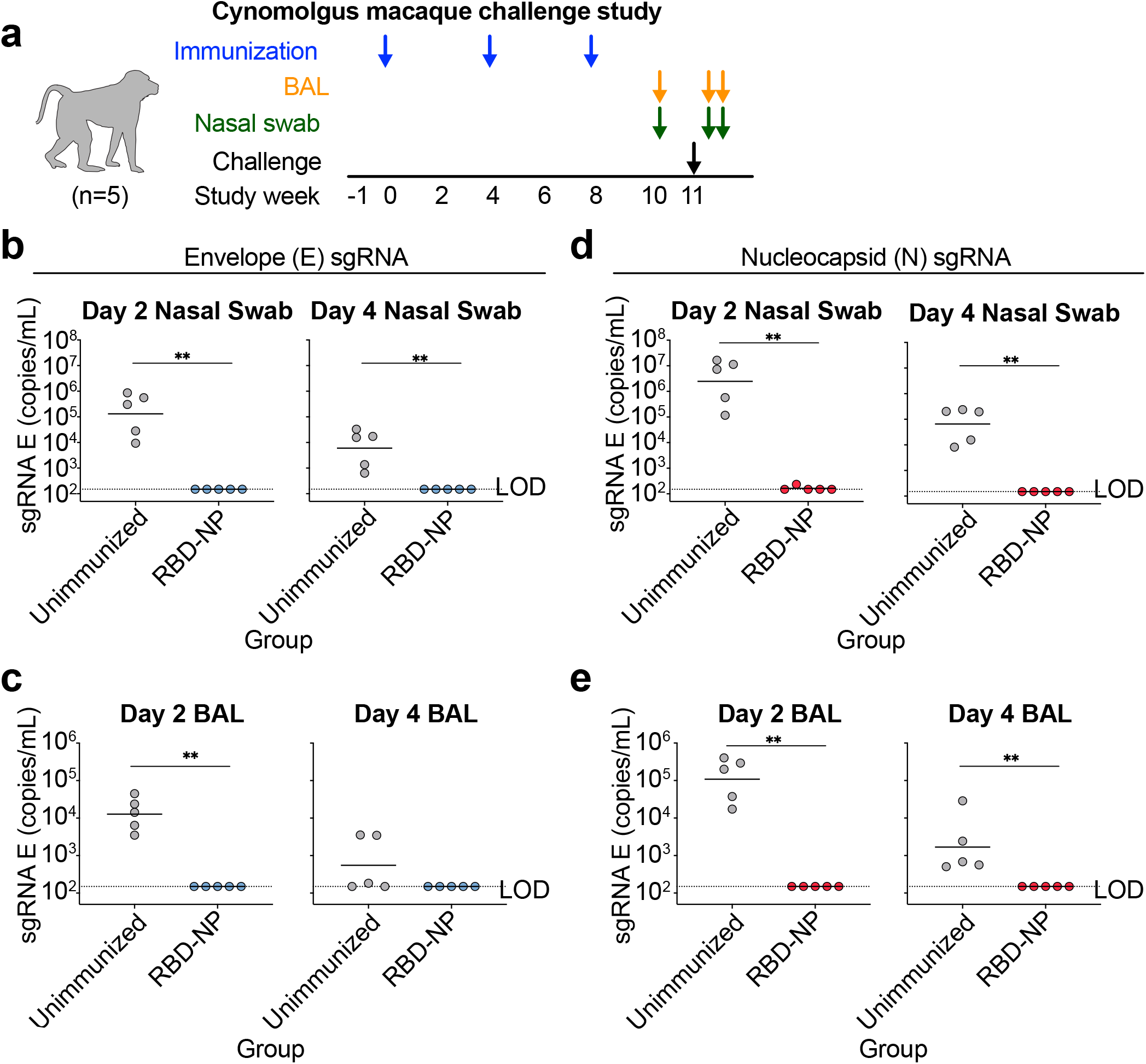
RBD-scNP vaccination completely prevents virus replication in the upper and lower respiratory tract after intranasal and intratracheal SARS-CoV-2 challenge in all but one macaque. **a.** Macaque intranasal/intratracheal SARS-CoV-2 challenge study design. Blue arrows indicate the time points for RBD-nanoparticle immunizations. Biospecimens were collected as indicated by the green and orange arrows. **b-e**. Quantification of viral subgenomic RNA (sgRNA) in unimmunized (gray symbols) and RBD-scNP-immunized (red or blue symbols) macaques. sgRNA encoding the **b, c** envelope (E) gene or **d, e** nucleocapsid (N) gene of SARS-CoV-2 was quantified two (left) and four (right) days after SARS-CoV-2 challenge. E or N gene sgRNA was measured in both **b, d** nasal swabs and **c, e** bronchoalveolar lavage (BAL) fluids from each macaque. Limit of detection (LOD) for the assay is 150 copies/mL. Each symbol represents an individual macaque with the group mean shown with a horizontal bar (\*\**P*<0.01, Two-tailed Exact Wilcoxon test n = 5).

This study demonstrates that immunization with SARS-CoV-2 Spike either as a protein RBD-scNP or as an mRNA-LNP elicits a cross-reactive antibody response capable of neutralizing multiple SARS-related human and bat betaCoVs. These results demonstrate that SARS-CoV-2 vaccination with either the RBD-scNP or the stabilized transmembrane spike mRNA-LNP vaccines currently approved for use in humans, will likely elicit cross-neutralizing antibodies with the potential to prevent future animal CoV spillover events to humans ^30,42,45^. The ~one log higher neutralization titers induced by RBD-scNP suggests improved durability of responses.

The identification of betaCoV cross-neutralizing antibodies such as DH1047 have shown that the SARS-CoV-2 RBD contains a conserved betaCoV group 2b cross-neutralizing epitope^13,23,24,46,47^. Vaccination of macaques in this study showed that cross-neutralizing epitopes on RBD can be potently targeted by primates. A recent study reported mosaic RBD nanoparticles arrayed with RBDs from various CoVs induced cross-neutralizing antibodies in mice ^30^. Here, we show that cross-neutralizing antibodies are not only elicited by RBD nanoparticles, but also by mRNA-LNPs expressing SARS-CoV-2 stabilized spike of the design currently in mRNA COVID-19 vaccines. The 24-mer RBD-scNP nanoparticle protein adjuvanted with a toll-like receptor agonist 3M-052 adsorbed to alum elicited the highest group 2b cross-neutralizing antibody titers. 3M-052 formulated with alum or in an emulsion has induced Iong-lasting memory B cells and protection against SHIV challenge in non-human primates^48,49^. A Phase I clinical study using 3M-052/Alum to induce neutralizing antibody responses to an HIV vaccine candidate is underway (NCT04177355). Thus, this vaccine modality represents a promising first-generation pan-group 2b betaCoV vaccine with the potential to durably inhibit future zoonotic transmission ^25^.

The emergence of SARS-CoV-2 neutralization-resistant and highly infectious variants continues to be a concern for vaccine efficacy. We found here that protein nanoparticle or mRNA-LNP SARS-CoV-2 spike immunization elicited SARS-CoV-2 neutralizing antibodies capable of neutralizing the predominant SARS-CoV-2 variant D614G as well as the newly-emerged B.1.1.7 UK variant. Thus, while the B.1.1.7 variant may be more transmissible, it was equally as sensitive to vaccine-induced serum neutralization as the predominant circulating SARS-CoV-2 D614G strain. Importantly, cross-reactive RBD-scNP immune sera and S-2P mRNA-LNP immune sera bound equally as well to spike ectodomain molecules with South African B1.351, Brazilian P.1 and UK B.1.1.7 RBD mutations, and are compatible with the recent demonstration that current COVID-19 vaccines have efficacy, albeit reduced, against the UK and South African SARS-CoV-2 variants^50–55^.

Protection against asymptomatic infection and the durability of protection remain concerns for current vaccines. The RBD-scNP vaccine induced robust protective immunity for SARS-CoV-2 replication in the upper and lower respiratory tract. This degree of virus suppression in the upper respiratory tract has not been routinely achieved with SARS-CoV-2 challenge in macaques ^56,57^ and supports the notion of inducing SARS-CoV-2 sterilizing immunity with vaccination. Additionally, the extraordinarily high neutralization titers achieved by RBD-scNP vaccination bode well for an extended duration of protection. Critically, as we have had 3 coronavirus epidemics in the past 20 years, there is a need to develop an effective pancoronavirus vaccine prior to the next pandemic. The RBD-scNP vaccine induces neutralizing antibodies to SARS-CoV-1, SARS-CoV-2, batCoV-WIV-1 and batCoV-SCH014 and represents a platform for producing a pancoronavirus vaccine that could prevent, rapidly temper, or extinguish the next spillover of a coronavirus into humans.

## Supporting information

Combined Supplemental information

## METHODS

### Animals and immunizations

Rhesus and cynomolgus macaques were housed and treated in AAALAC-accredited institutions. The study protocol and all veterinarian procedures were approved by the Bioqual IACUC per a memorandum of understanding with the Duke IACUC, and were performed based on standard operating procedures. Nucleoside-modified messenger RNA encapsulated in lipid nanoparticles (mRNA-LNP) was prepared as previously stated ^58,59^. Rhesus macaques (n=8) were immunized intramuscularly with 50μg of mRNA-LNP encoding the receptor binding domain monomer. Cynomolgus macaques (n=5) were immunized with either 50μg of mRNA-LNP encoding the transmembrane Spike protein stabilized with K986P and V987P mutations. An additional group of cynomolgus macaques (n=5) were immunized in the quadriceps with 100 μg of RBD-scNP adjuvanted with 5 μg of 3M-052 formulated in alum^33^. Biospecimens before challenge, 2 days post-challenge, and 4 days post challenge were collected as described previously ^24^.

### SARS-CoV-2 intranasal and intratracheal challenge

All animals were challenged at week 11 (3 weeks after last vaccination) through combined intratracheal (IT, 3.0 mL) and intranasal (IN, 0.5 mL per nostril) inoculation with an infectious dose of 105 PFU of SARS-CoV-2 (2019-nCoV/USA-WA1/2020). The stock was generated at BIOQUAL (lot# 030120-1030, 3.31 × 10^5^ PFU/mL) from a p4 seed stock obtained from BEI Resources (NR-52281). The stock underwent deep sequencing to confirm homology with the WA1/2020 isolate. Virus was stored at −80°C prior to use, thawed by hand and placed immediately on wet ice. Stock was diluted to 2.5×10^4^ PFU/mL in PBS and vortexed gently for 5 seconds prior to inoculation. Nasal swabs, bronchoalveolar lavage (BAL), plasma, and serum samples were collected seven days before, two days after, and four days after challenge. Unimmunized macaques (n=50 were used as a negative control group. Protection from SARS-CoV-2 infection was determined by quantitative PCR of SARS-CoV-2 subgenomic envelope (E) and the more sensitive nucleocapsid (N) RNA (E or N sgRNA) ^39^ as stated below.

### SARS-CoV-2 protein production

The CoV ectodomain constructs were produce and purified as describe previously ^60^. The Spike (S) ectodomain was stabilized by the introduction of 2 prolines at amino acid positions 986 and 987 and referred to as S-2P. Plasmids encoding Spike-2P and HexaPro ^61^ were transiently transfected in FreeStyle 293 cells (Thermo Fisher) using Turbo293 (SpeedBiosystems) or 293Fectin (ThermoFisher). The constructs contained an HRV 3C-cleavable C-terminal twinStrepTagII-8xHis tag. On day 6, cell-free culture supernatant was generated by centrifugation of the culture and filtering through a 0.8 um filter. Protein was purified from filtered cell culture supernatants by StrepTactin resin (IBA) and by size exclusion chromatography using Superose 6 column (GE Healthcare) in 10 mM Tris pH8,150 mM NaCl or 2 mM Tris pH 8, 200 mM NaCl, 0.02% NaN_3_. ACE-2-Fc was expressed by transient transfection of Freestyle 293-F cells^60^. ACE2-Fc was purified from cell culture supernatant by HiTrap protein A column chromatography and Superdex200 size exclusion chromatography in 10 mM Tris pH8,150 mM NaCl. SARS-CoV-2 NTD was produced as previously described ^62^. SARS-CoV-2 fusion peptide was synthesized (GenScript).

### Sortase A conjugation of SARS-CoV-2 RBD to *H. pylori* ferritin nanoparticles

Wuhan strain SARS-CoV-2 RBD was expressed with sortase A donor sequence LPETGG encoded at its c-terminus. C-terminal to the sortase A donor sequence was an HRV-3C cleavage site, 8× his tag, and a twin StrepTagII (IBA). The SARS-CoV-2 RBD was expressed in Freestyle293 cells and purified by StrepTactin affinity chromatography (IBA) and superdex200 size exclusion chromatography as stated above. *H. pylori* ferritin particles were expressed with a pentaglycine sortase A acceptor sequence encoded at its N-terminus of each subunit. For affinity purification of ferritin particles, 6XHis tags were appended C-terminal to a HRV3C cleavage site. Ferritin particles with a sortase A N-terminal tag were buffer exchanged into 50mM Tris, 150mM NaCI, 5mM CaCI2, pH7.5. 180 μM SARS-CoV-2 RBD was mixed with 120 μM of ferritin subunits and incubated with 100 μM of sortase A overnight at room temperature. Following incubation conjugated particles were isolated from free ferritin or free RBD by size exclusion chromatography using a Superose6 16/60 column.

### Biolayer interferometry binding assays

Binding was measured using an OctetRed 96 (ForteBio). Anti-human IgG capture (AHC) sensor tips (Forte Bio) were hydrated for at least 10 minutes in PBS. ACE2 and monoclonal antibodies were diluted to 20 μg/mL in PBS and placed in black 96-well assay plate. The influenza antibody CH65 was used as the background reference antibody. The RBD nanoparticle was diluted to 50 μg/mL in PBS and added to the assay plate. Sensor tips were loaded with antibody for 120 s. Subsequently, the sensor tips were washed for 60 s in PBS to removed unbound antibody. The sensor tips were incubated in a fresh well of PBS to establish baseline reading before being dipped into RBD-scNP to allow association for 400 s. To measure dissociation of the antibody-RBD-scNP complex, the tip was incubated in PBS for 600 s. At the end of dissociation, the tip was ejected and a new tip was attached to load another antibody. The data was analyzed with Data Analysis HT v12 (ForteBio). Background binding observed with CH65 was subtracted from all values. All binding curves were aligned to the start of association. The binding response at the end of the 400 s association phase was plotted in GraphPad Prism v9.0.

### Surface plasmon resonance (SPR) assays

SPR measurements of DH1047 antigen binding fragment (Fab) binding to monomeric SARS-CoV-2 receptor binding domain (RBD) proteins were performed in HBS-EP+ running buffer using a Biacore S200 instrument (Cytiva). Assays were performed in the DHVI BIA Core Facility. The RBD was first captured via its twin-StrepTagII onto a Series S Streptavidin chip to a level of 300-400 resonance units (RU). The antibody Fabs were injected at 0.5 to 500 nM over the captured S proteins using the single cycle kinetics injection mode at a flow rate of 50 μL/min. Fab association occurred for 180 s followed by a dissociation of 360 seconds after the end of the association phase. At the end of the dissociation phase the RBD was regenerated with a 30 s injection of glycine pH1.5. Binding values were analyzed with Biacore S200 Evaluation software (Cytiva). References included blank streptavidin surface along with blank buffer binding and was subtracted from DH1047 values to account for signal drift and non-specific protein binding. A 1:1 Langmuir model with a local Rmax was used for curve fitting. Binding rates and constants were derived from the curve. Representative results from two independent experiments are shown.

### SARS-CoV-2 pseudovirus neutralization

Neutralization of SARS-CoV-2 Spike-pseudotyped virus was performed by adapting an infection assay described previously with lentiviral vectors and infection in 293T/ACE2.MF (the cell line was kindly provided by Drs. Mike Farzan and Huihui Mu at Scripps). Cells were maintained in DMEM containing 10% FBS and 50 μg/ml gentamicin. An expression plasmid encoding codon-optimized full-length spike of the Wuhan-1 strain (VRC7480), was provided by Drs. Barney Graham and Kizzmekia Corbett at the Vaccine Research Center, National Institutes of Health (USA). The D614G mutation was introduced into VRC7480 by site-directed mutagenesis using the QuikChange Lightning Site-Directed Mutagenesis Kit from Agilent Technologies (Catalog # 210518). The mutation was confirmed by full-length spike gene sequencing. Pseudovirions were produced in HEK 293T/17 cells (ATCC cat. no. CRL-11268) by transfection using Fugene 6 (Promega, Catalog #E2692). Pseudovirions for 293T/ACE2 infection were produced by co-transfection with a lentiviral backbone (pCMV ΔR8.2) and firefly luciferase reporter gene (pHR’ CMV Luc) ^63^. Culture supernatants from transfections were clarified of cells by low-speed centrifugation and filtration (0.45 μm filter) and stored in 1 ml aliquots at −80 °C.

For 293T/ACE2 neutralization assays, a pre-titrated dose of virus was incubated with 8 serial 3-fold or 5-fold dilutions of mAbs in duplicate in a total volume of 150 μL for 1 hr at 37 °C in 96-well flat-bottom poly-L-lysine-coated culture plates (Corning Biocoat). Cells were suspended using TrypLE express enzyme solution (Thermo Fisher Scientific) and immediately added to all wells (10,000 cells in 100 μL of growth medium per well). One set of 8 control wells received cells + virus (virus control) and another set of 8 wells received cells only (background control). After 66-72 h of incubation, medium was removed by gentle aspiration and 30 μL of Promega 1× lysis buffer was added to all wells. After a 10-minute incubation at room temperature, 100 μl of Bright-Glo luciferase reagent was added to all wells. After 1-2 minutes, 110 μl of the cell lysate was transferred to a black/white plate (Perkin-Elmer). Luminescence was measured using a PerkinElmer Life Sciences, Model Victor2 luminometer. Neutralization titers are the serum dilution (ID50/ID80) at which relative luminescence units (RLU) were reduced by 50% and 80% compared to virus control wells after subtraction of background RLUs.

### Plasma IgG blocking of ACE2 binding

For ACE2 blocking assays, plates were coated with 2 μg/mL recombinant ACE2 protein, then washed and blocked with 3% BSA in 1× PBS. While assay plates blocked, purified antibodies were diluted as stated above, only in 1% BSA with 0.05% Tween-20. In a separate dilution plate Spike-2P protein was mixed with the antibodies at a final concentration equal to the EC50 at which spike binds to ACE2 protein. The mixture was allowed to incubate at room temperature for 1 hour. Blocked assay plates were then washed and the antibody-spike mixture was added to the assay plates for a period of 1 hour at room temperature. Plates were washed and a polyclonal rabbit serum against the same spike protein (nCoV-1 nCoV-2P.293F) was added for 1 hour, washed and detected with goat anti rabbit-HRP (Abcam cat# ab97080) followed by TMB substrate. The extent to which antibodies were able to block the binding spike protein to ACE2 was determined by comparing the OD of antibody samples at 450 nm to the OD of samples containing spike protein only with no antibody. The following formula was used to calculate percent blocking: blocking% = (100 - (OD sample/OD of spike only)*100).

### Plasma IgG blocking of RBD monoclonal antibody binding

Blocking assays for DH1041 and DH1047 were performed as stated above for ACE2, except plates were coated with either DH1041 or DH1047 instead of ACE2.

### Plasma IgG ELISA binding assays

For ELISA binding assays of Coronavirus Spike antibodies, the antigen panel included SARS-CoV-2 Spike S1+S2 ectodomain (ECD) (SINO, Catalog # 40589-V08B1), SARS-CoV-2 Spike-2P ^60^, SARS-CoV-2 Spike S2 ECD (SINO, Catalog # 40590-V08B), SARS-CoV-2 Spike RBD from insect cell sf9 (SINO, Catalog # 40592-V08B), SARS-CoV-2 Spike RBD from mammalian cell 293 (SINO, Catalog # 40592-V08H), SARS-CoV-2 Spike NTD-Biotin, SARS-CoV Spike Protein DeltaTM (BEI, Catalog # NR-722), SARS-CoV WH20 Spike RBD (SINO, Catalog # 40150-V08B2), SARS-CoV WH20 Spike S1 (SINO, Catalog #40150-V08B1), SARS-CoV-1 RBD, MERS-CoV Spike S1+S2 (SINO, Catalog # 40069-V08B), MERS-CoV Spike S1 (SINO, Catalog #40069-V08B1), MERS-CoV Spike S2 (SINO, Catalog #40070-V08B), MERS-CoV Spike RBD (SINO, Catalog #40071-V08B1), MERS-CoV Spike RBD.

For binding ELISA, 384-well ELISA plates were coated with 2 μg/mL of antigens in 0.1 M sodium bicarbonate overnight at 4°C. Plates were washed with PBS + 0.05% Tween 20 and blocked with assay diluent (PBS containing 4% (w/v) whey protein, 15% Normal Goat Serum, 0.5% Tween-20, and 0.05% Sodium Azide) at room temperature for 1 hour. Purified mAb samples in 3-fold serial dilutions in assay diluent starting at 100 μg/mL were added and incubated for 1 h, followed by washes with PBS-0.1% Tween 20. HRP-conjugated goat anti-human IgG secondary Ab (SouthernBiotech, catalog #2040-05) was diluted to 1:10,000 and incubated at room temperature for 1 hour. These plates were washed four times and developed with tetramethylbenzidine substrate (SureBlue Reserve-KPL). The reaction was stopped with 1 M HCl, and optical density at 450 nm (OD_450_) was determined.

### Subgenomic RNA real time PCR quantification

The assay for SARS-CoV-2 quantitative Polymerase Chain Reaction (qPCR) detects total RNA using the WHO primer/probe set E_Sarbeco (Charité/Berlin). A QIAsymphony SP (Qiagen, Hilden, Germany) automated sample preparation platform along with a virus/pathogen DSP midi kit and the *complex800* protocol were used to extract viral RNA from 800 μL of pooled samples. A reverse primer specific to the envelope gene of SARS-CoV-2 (5’-ATA TTG CAG CAG TAC GCA CAC A-3’) was annealed to the extracted RNA and then reverse transcribed into cDNA using SuperScript™ III Reverse Transcriptase (Thermo Fisher Scientific, Waltham, MA) along with RNAse Out (Thermo Fisher Scientific, Waltham, MA). The resulting cDNA was treated with RNase H (Thermo Fisher Scientific, Waltham, MA) and then added to a custom 4x TaqMan™ Gene Expression Master Mix (Thermo Fisher Scientific, Waltham, MA) containing primers and a fluorescently labeled hydrolysis probe specific for the envelope gene of SARS-CoV-2 (forward primer 5’-ACA GGT ACG TTA ATA GTT AAT AGC GT-3’, reverse primer 5’-ATA TTG CAG CAG TAC GCA CAC A-3’, probe 5’-6FAM/AC ACT AGC C/ZEN/A TCC TTA CTG CGC TTC G/IABkFQ-3’). The qPCR was carried out on a QuantStudio 3 Real-Time PCR System (Thermo Fisher Scientific, Waltham, MA) using the following thermal cycler parameters: heat to 50°C, hold for 2 min, heat to 95°C, hold for 10 min, then the following parameters are repeated for 50 cycles: heat to 95°C, hold for 15 seconds, cool to 60°C and hold for 1 minute. SARS-CoV-2 RNA copies per reaction were interpolated using quantification cycle data and a serial dilution of a highly characterized custom DNA plasmid containing the SARS-CoV-2 envelope gene sequence. Mean RNA copies per milliliter were then calculated by applying the assay dilution factor (DF=11.7). The limit of detection (LOD) for this assay is approximately 62 RNA copies per mL of sample.

### Live virus neutralization assays

Full-length SARS-CoV-2, SARS-CoV, WIV-1, and RsSHC014 viruses were designed to express nanoluciferase (nLuc) and were recovered via reverse genetics as described previously ^64–66^. Virus titers were measured in Vero E6 USAMRIID cells, as defined by plaque forming units (PFU) per ml, in a 6-well plate format in quadruplicate biological replicates for accuracy. For the 96-well neutralization assay, Vero E6 USAMRID cells were plated at 20,000 cells per well the day prior in clear bottom black walled plates. Cells were inspected to ensure confluency on the day of assay. Serum samples were tested at a starting dilution of 1:20 and were serially diluted 3-fold up to nine dilution spots. Serially diluted serum samples were mixed in equal volume with diluted virus. Antibody-virus and virus only mixtures were then incubated at 37°C with 5% CO_2_ for one hour. Following incubation, serially diluted sera and virus only controls were added in duplicate to the cells at 75 PFU at 37°C with 5% CO_2_. After 24 hours, cells were lysed, and luciferase activity was measured via Nano-Glo Luciferase Assay System (Promega) according to the manufacturer specifications. Luminescence was measured by a Spectramax M3 plate reader (Molecular Devices, San Jose, CA). Virus neutralization titers were defined as the sample dilution at which a 50% reduction in RLU was observed relative to the average of the virus control wells.

### Biocontainment and biosafety

All work described here was performed with approved standard operating procedures for SARS-CoV-2 in a biosafety level 3 (BSL-3) facility conforming to requirements recommended in the Microbiological and Biomedical Laboratories, by the U.S. Department of Health and Human Service, the U.S. Public Health Service, and the U.S. Center for Disease Control and Prevention (CDC), and the National Institutes of Health (NIH).

### Recombinant IgG production

Expi293-F cells were diluted to 2.5E6 cells/mL on the day of transfection. Cells were co-transfected with Expifectamine and heavy and light chain expression plasmids. Enhancers were added 16h after transfection. On day 5, the cell culture was cleared of cells by centrifugation, filtered, and incubated with protein A beads overnight. The next day the protein A resin was washed with Tris buffered saline and then added to a 25 mL column. The resin was washed again and then glacial acetic acid was used to elute antibody off of the protein A resin. The pH of the solution was neutralized with 1M Tris pH8. The antibody was buffer exchanged into 25 mM sodium citrate pH6 supplemented with 150 mM NaCl, 0.2 μm filtered, and frozen at −80°C.

### Negative stain electron microscopy

The RBD nanoparticle protein at ~1-5 mg/ml concentration that had been flash frozen and stored at −80 °C was thawed in an aluminum block at 37 °C for 5 minutes; then 1-4 μL of RBD nanoparticle was diluted to a final concentration of 0.1 mg/ml into room-temperature buffer containing 150 mM NaCl, 20 mM HEPES pH 7.4, 5% glycerol, and 7.5 mM glutaraldehyde. After 5 minutes cross-linking, excess glutaraldehyde was quenched by adding sufficient 1 M Tris pH 7.4 stock to give a final concentration of 75 mM Tris and incubated for 5 minutes. For negative stain, carbon-coated grids (EMS, CF300-cu-UL) were glow-discharged for 20s at 15 mA, after which a 5-μl drop of quenched sample was incubated on the grid for 10-15 s, blotted, and then stained with 2% uranyl formate. After air drying grids were imaged with a Philips EM420 electron microscope operated at 120 kV, at 82,000× magnification and images captured with a 2k × 2k CCD camera at a pixel size of 4.02 Å.

### Processing of negative stain images

The RELION 3.0 program was used for all negative stain image processing. Images were imported, CTF-corrected with CTFFIND, and particles were picked using a spike template from previous 2D class averages of spike alone. Extracted particle stacks were subjected to 2-3 rounds of 2D class averaging and selection to discard junk particles and background picks. Cleaned particle stacks were then subjected to 3D classification using a starting model created from a bare spike model, PDB 6vsb, low-pass filtered to 30 Å. Classes that showed clearly-defined Fabs were selected for final refinements followed by automatic filtering and B-factor sharpening with the default Relion post-processing parameters.

### Betacoronavirus sequence analysis

Heatmaps of amino acid sequence similarity were computed for a representative set of betacoronaviruses using the ComplexHeatmap package in R. Briefly, 1408 betacoronavirus sequences were retrieved from NCBI Genbank, aligned to the Wuhan-1 spike protein sequence, and trimmed to the aligned region. The 1408 spike sequences were then clustered using USEARCH ^67^ with a sequence identity threshold of 0.90 resulting in 52 clusters. We sampled one sequence from each cluster to generate a representative set of sequences. Five betacoronavirus sequences of interest not originally included in the clustered set were added: SARS-CoV-2, GXP4L, batCoV-RaTG13, batCoV-SHC014, batCoV-WIV-1. This resulted in a set of 57 representative spike sequences. Pairs of spike amino acid sequences were aligned using a global alignment and the BLOSUM62 scoring matrix. For RBD and NTD domain alignments, spike sequences were aligned to the Wuhan 1 spike protein RBD region (residues 330-521) and NTD region (residues 27-292), respectively, and trimmed to the aligned region. Phylogenetic tree construction of RBD sequences was performed with Geneious Prime 2020.1.2 using the Neighbor Joining method and default parameters. To map group 2b betaCoV sequence conservation onto the RBD structure, group 2b spike sequences were retrieved from Genbank and clustered using USEARCH ^67^ with a sequence identity threshold of 0.99 resulting in 39 clusters. For clusters of size >5, 5 spike sequences were randomly downsampled from each cluster. The resulting set of 73 sequences was aligned using MAFFT ^68^. Conservation scores for each position in the multiple sequence alignment were calculated using the trident scoring method ^69^ and computed using the MstatX program (https://github.com/gcollet/MstatX). The conservation scores were then mapped to the RBD domain coordinates (PDB: 7LD1) and images rendered with PyMol version 2.3.5.

### Statistics Analysis

Data were plotted using Prism GraphPad 8.0. Wilcoxon rank sum exact test was performed to compare differences between groups with p-value < 0.05 considered significant using SAS 9.4 (SAS Institute, Cary, NC). No adjustments were made to the p-values for multiple comparisons. IC50 and IC80 values were calculated using R statistical software (version 4.0.0; R Foundation for Statistical Computing, Vienna, Austria). The R package ‘nplr’ was used to fit 4-Parameter Logistic (4-PL) regression curves to the average values from duplicate experiments, and these fits were used to estimate the concentrations corresponding to 50% and 80% neutralization.

## ACKNOWLEDGEMENTS

We thank Victoria Gee-Lai, Margaret Deyton, Charlene McDanal, Brian Watts, and Kenneth Cronin for technical assistance. We thank Elizabeth Donahue for program management and assistance with manuscript preparation. We thank John Harrison, Alex Granados, Adrienne Goode, Anthony Cook, Alan Dodson, Katelyn Steingrebe, Bridget Bart, Laurent Pessaint, Alex VanRy, Daniel Valentin, Amanda Strasbaugh, and Mehtap Cabus for assistance with macaque studies. We thank Shelby O’Connor and John James Baczenas at the Dept. of Pathology & Laboratory Medicine, University of Wisconsin-Madison for sequencing support. The following reagent was deposited by the Centers for Disease Control and Prevention and obtained through BEI Resources, NIAID, NIH: SARS-Related Coronavirus 2, Isolate USA-WA1/2020, NR-52281. Supported by a grant from the State of North Carolina with funds from the federal CARES Act, by fund from NIH, NIAID, DAIDS grant AI142596, and by R01AI157155 (RSB) and U54 CA260543 (RSB). This project was also supported by the North Carolina Policy Collaboratory at the University of North Carolina at Chapel Hill with funding from the North Carolina Coronavirus Relief Fund established and appropriated by the North Carolina General Assembly. This study was also supported by funding from an NIH F32 AI152296, a Burroughs Wellcome Fund Postdoctoral Enrichment Program Award, and was previously supported by an NIH NIAID T32 AI007151 (all three awarded to DRM). COVID sample processing was performed in the Duke Regional Biocontainment Laboratory, which received partial support for construction from the NIH/NIAD (UC6AI058607; GDS) with support from a cooperative agreement with DOD/DARPA (HR0011-17-2-0069; GDS).

## AUTHOR CONTRIBUTIONS

KOS and BFH designed and managed the study, reviewed all data and wrote and edited the manuscript; DW, NP designed and produced the mRNA-LNPs; EL, AMS, FPC, and HC expressed proteins; EL, RP, MB, THO, DRM, DCM, LVT, TDS, RSB carried out binding or neutralization assays; RSB and DRM prepared recombinant live viruses encoding nLuc; DL, CTM, TD, MG, DCD designed or performed sg or genomic RNA assays; RJE, SG, PA, KatM, KarM, MA, MB, and KW performed structural or sequence analysis; SMA performed SPR; LLS, MGL, HA, RS and IM performed or evaluated monkey studies; CWW, EWP, GDS collected, annotated, COVID-19 samples; MT selected, provided adjuvant; RWR and RLS performed statistical analyses; All authors edited and approved the manuscript.

## Supplementary information is available for this paper

**Correspondence** and requests for materials should be addressed to

**Reprints and permissions information is available at** http://www.nature.com/reprints

## Competing financial interests

MAT is an employee of 3M Company. 3M company had no role in the execution of the study, data collection, or data interpretation. BFH, KOS, and MAT have filed patents regarding the nanoparticle vaccine. DW is named on patents that describe the use of nucleoside-modified mRNA as a platform to deliver therapeutic proteins. DW and NP are also named on a patent describing the use of nucleoside-modified mRNA in lipid nanoparticles as a vaccine platform. All other authors declare no competing interests.

