## Supplementary material for "SARS-CoV-2 vaccination induces neutralizing antibodies against pandemic and pre-emergent SARS-related coronaviruses in monkeys": Combined Supplemental information

### Extended Data Figure 1

**a**

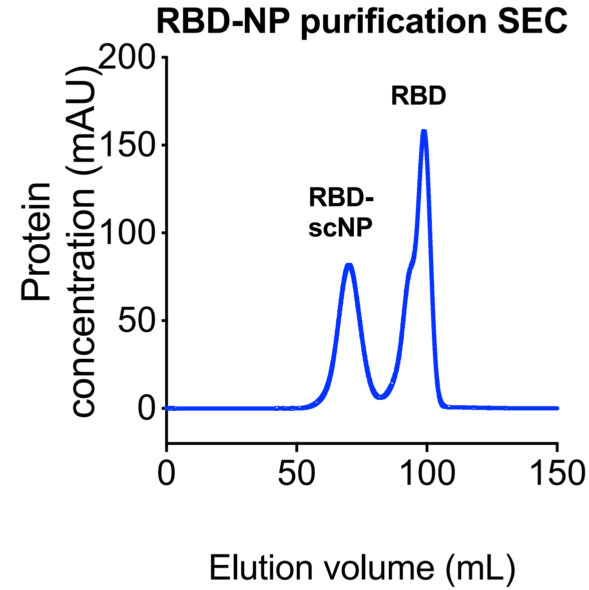


**b**

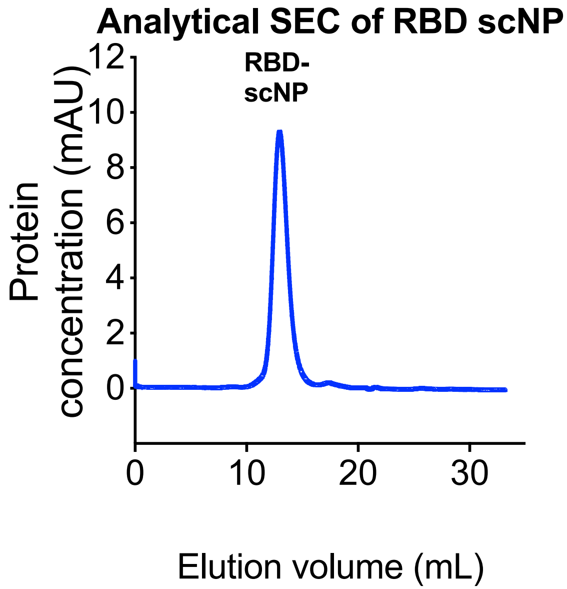


**c**

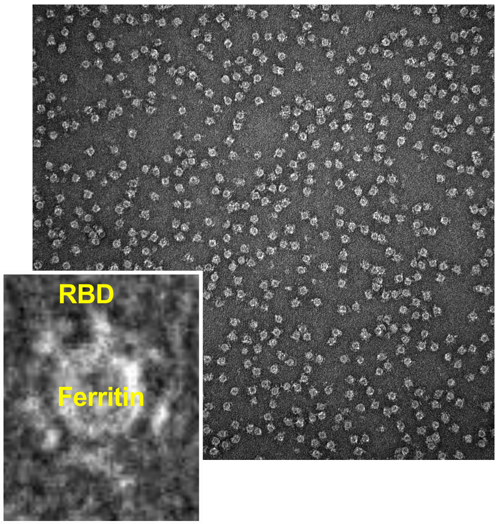


**d**

3M-052

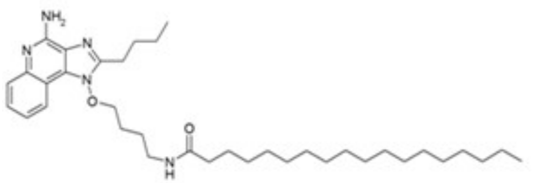

Molecular Weight = 593.90
Exact mass = 593
Molecular formula = C36H59N5O2
Molecular composition = C 72.81%, H 10.01%, N 11.79%, O 5.39%

**Extended Data Figure 1. Molecular and structural characterization of the SARS-CoV-2 RBD sortase A conjugated nanoparticle.**

**a** Size exclusion chromatography of RBD and ferritin sortase conjugation. The first peak shows conjugated protein. The second peak contains unconjugated RBD.

**b** Analytical size exclusion trace shows a homogenous nanoparticle preparation.

**c** Negative stain electron microscopy image of RBD-scNPs on a carbon grid. Inset shows a zoomed image of RBD-scNP. The zoomed image shows RBD molecules arrayed around the outside of the ferritin nanoparticle.

**d** Chemical structure of toll-like receptor 7 and 8 agonist 3M-052. Alum formulation of 3M-052 was used to adjuvant RBD-scNP immunization.

**Extended Data Figure 2**

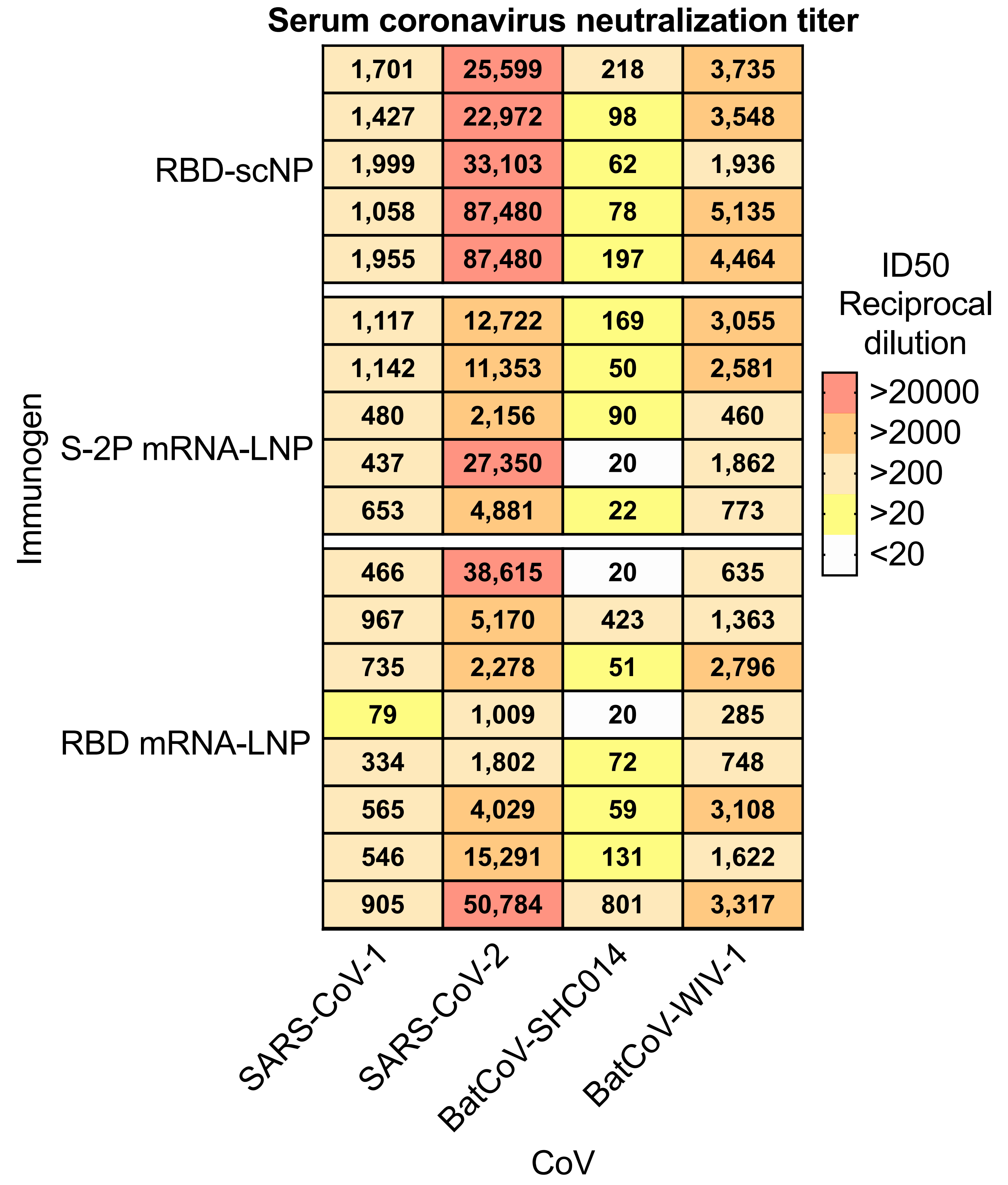

**Extended data figure 2. Cross-neutralizing antibodies are elicited by recombinant protein RBD-scNP and Spike mRNA-LNP immunization.** Each row shows neutralization titer for an individual macaque immunized with one of the three immunogens. Reciprocal serum dilutions titers of 87,480 is the upper limit of detection and 20 is lower limit of detection for this assay. Titers are derived from a nonlinear regression curve fitted to the average of duplicate measurements.

### Extended Data Figure 3

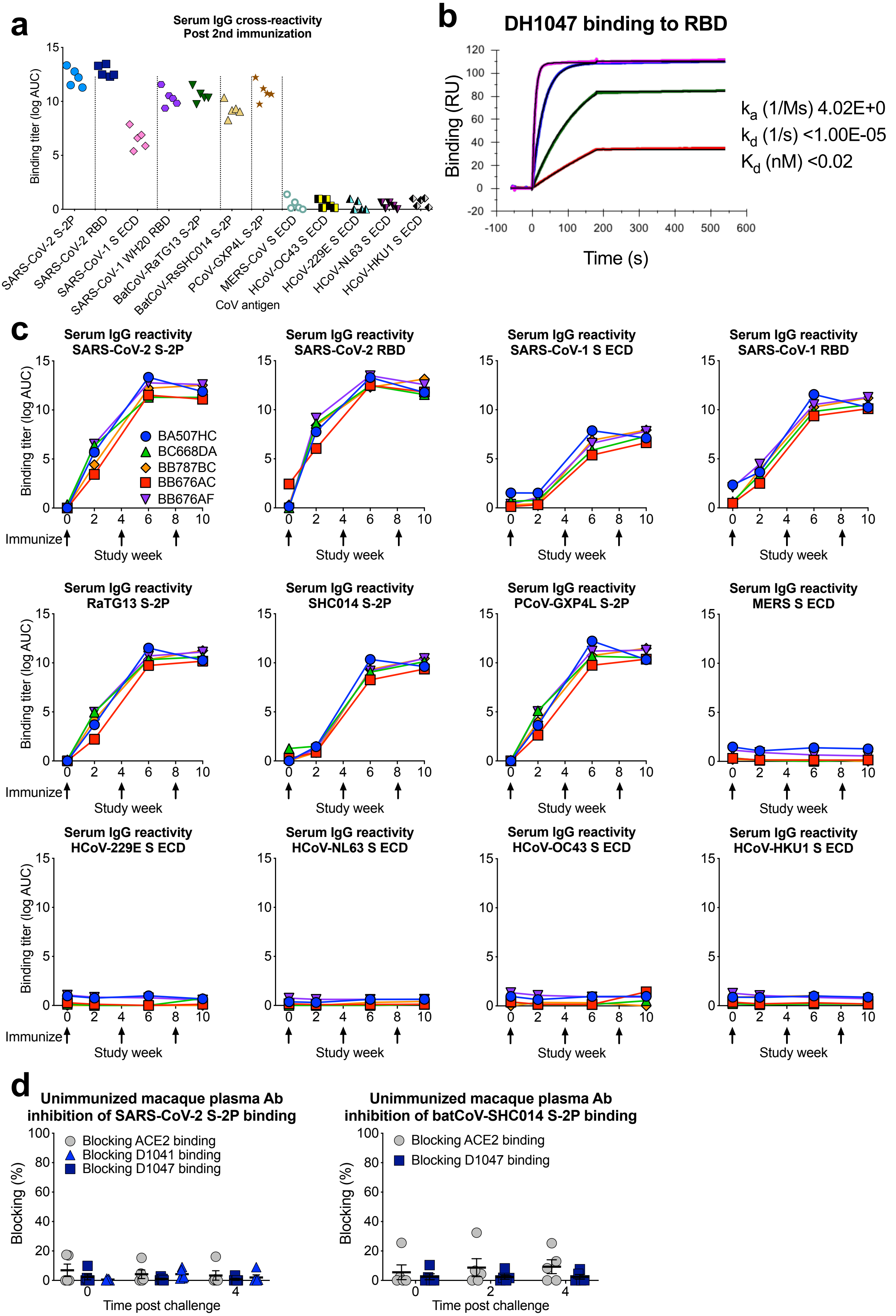

**Extended Data Figure 3. Cross-reactive plasma antibody responses elicited by RBD-NP immunization in macaques.**

**a** Plasma IgG from macaques immunized twice with RBD-scNP binds to Spike from human, bat, and pangolin SARS-related coronavirus Spike (S) in ELISA, but not endemic human coronaviruses or MERS-CoV. ECD, ectodomain.

**b** Determination of DH1047 antigen binding fragment (Fab) binding kinetics to RBD monomer by surface plasmon resonance. Each curve shows a different concentration of DH1047 Fab. Binding kinetics are shown to the right from a 1:1 model fit.

**c** Time course of vaccinated macaque plasma IgG binding to human, bat, and pangolin coronavirus S protein by ELISA. Each curve indicates the binding titer for an individual macaque. Arrows indicate immunization time points.

**d** Unimmunized macaque plasma antibody blocking of SARS-CoV-2 S-2P (left) and batCoV-SHC014 (right) binding to ACE2-Fc, RBD neutralizing antibody DH1041, and RBD cross-neutralizing antibody DH1047. Each symbol represents an individual macaque. Black horizontal bars indicate group mean and standard error.

### Extended data Figure 4

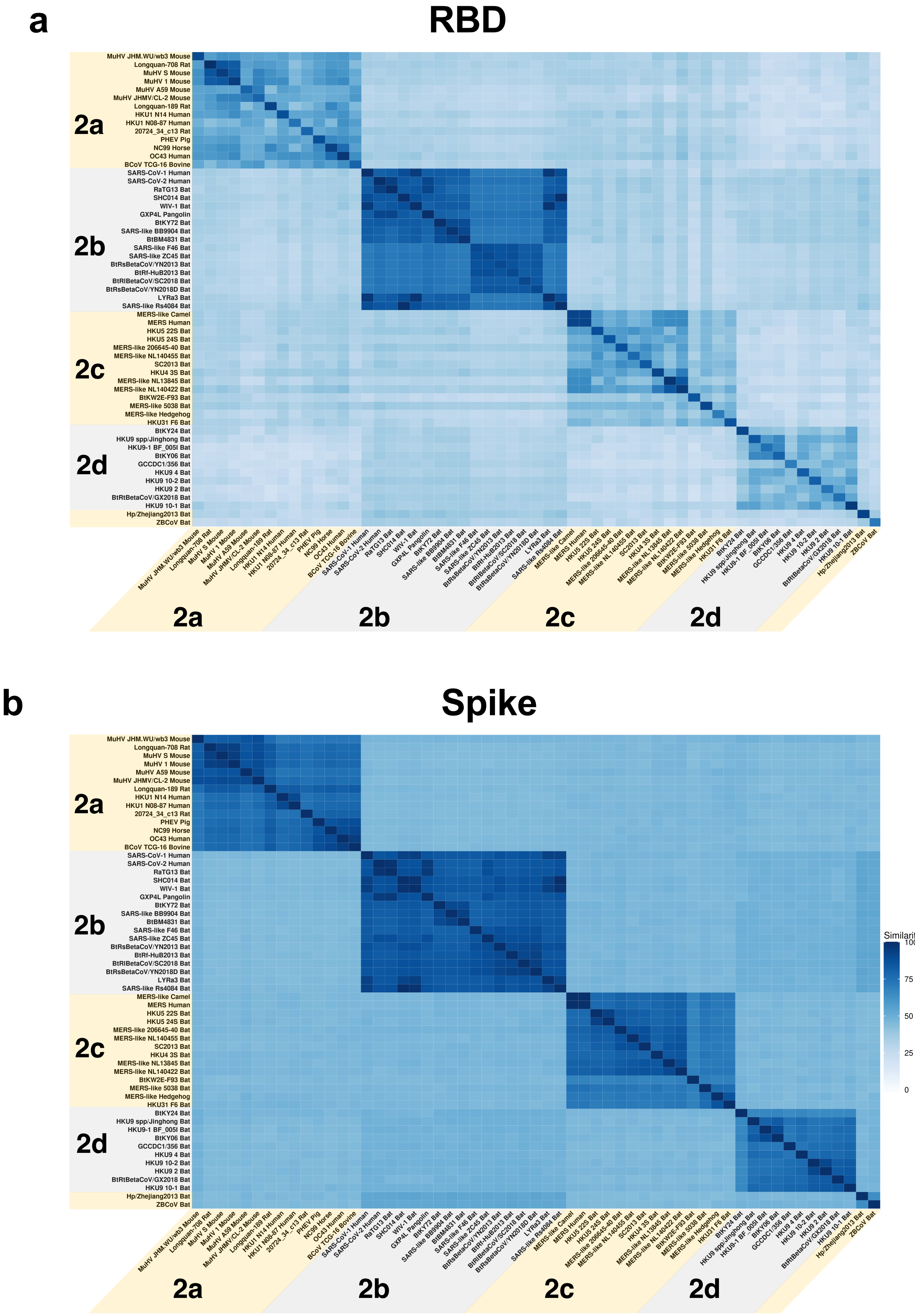

**Extended Data Figure 4. Sequence conservation among SARS-related betaCoV, MERS-CoV, and endemic human CoVs.**

**a,b.** Sequence similarity of **a** RBD and **b** spike protein for representative betacoronaviruses. Heatmaps displaying pairwise amino acid sequence similarity for 57 representative betacoronaviruses. Dark blue shading indicates high sequence similarity.

### Extended Data Figure 5

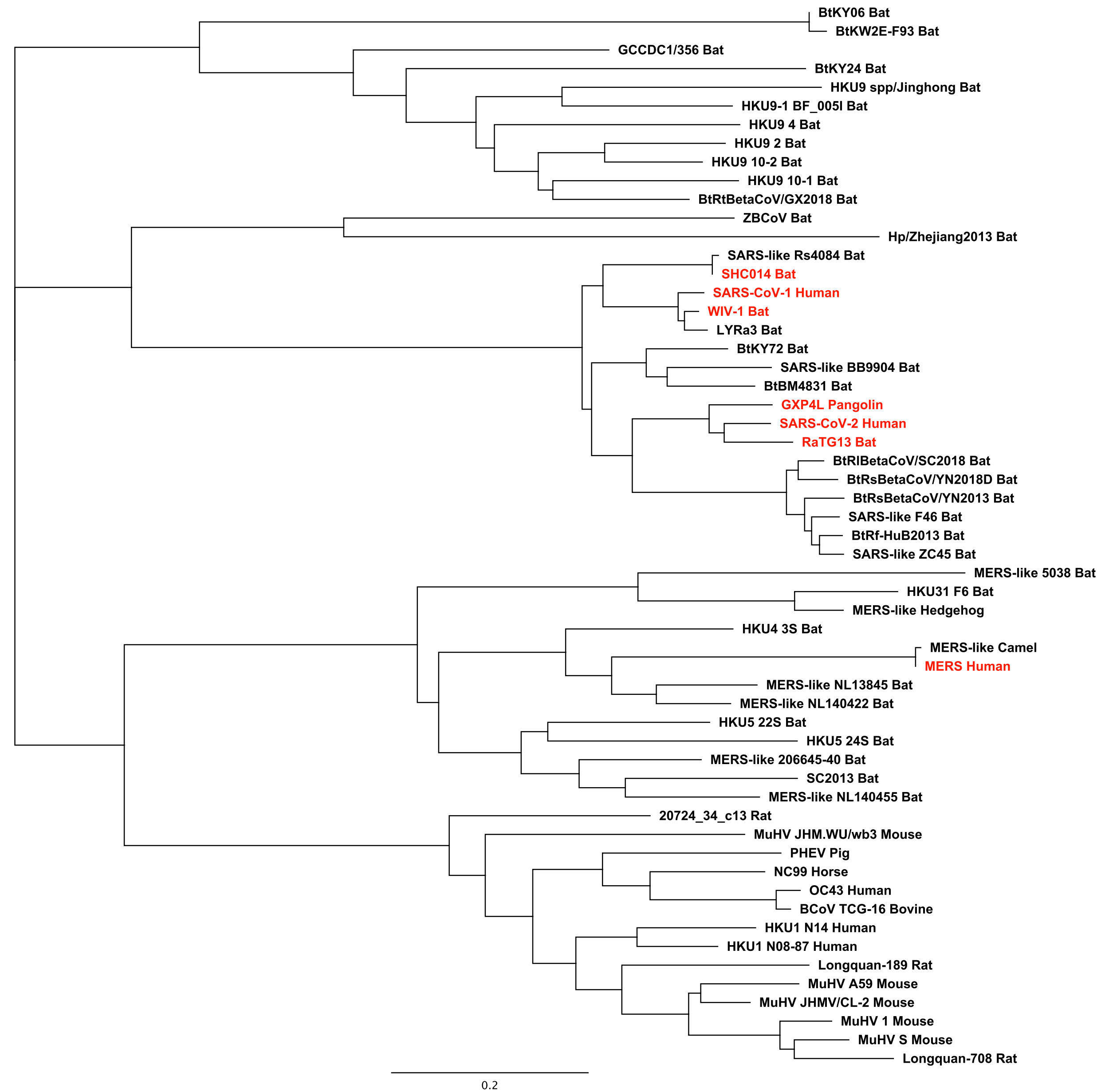

**Extended data figure 5**.. **Phylogenetic tree of representative betacoronavirus RBD sequences.** Group 2b betaCoVs of interest are shown highlighted in red. Branch length units are substitutions per site.

### Extended Data Figure 6

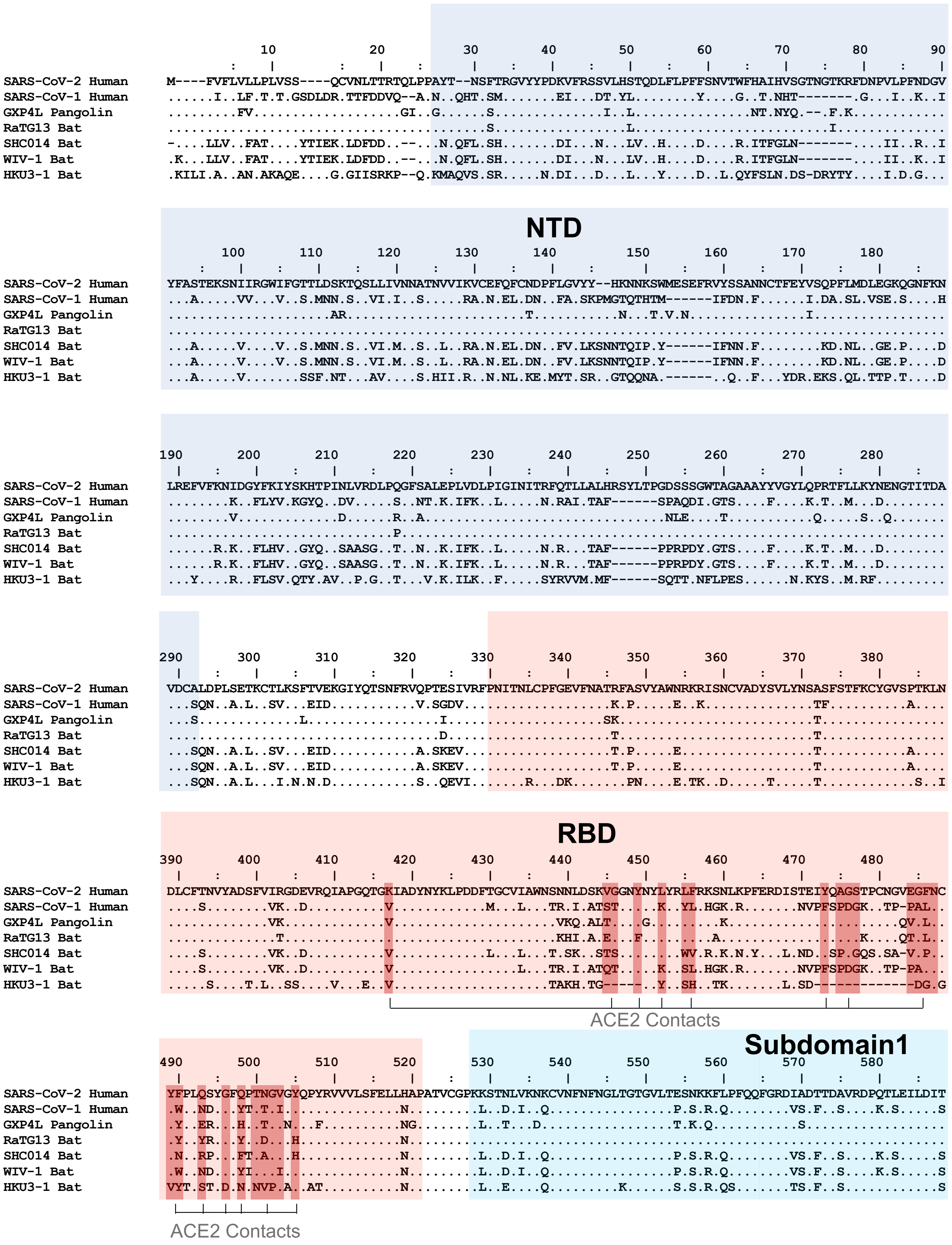

### Extended data Figure 6

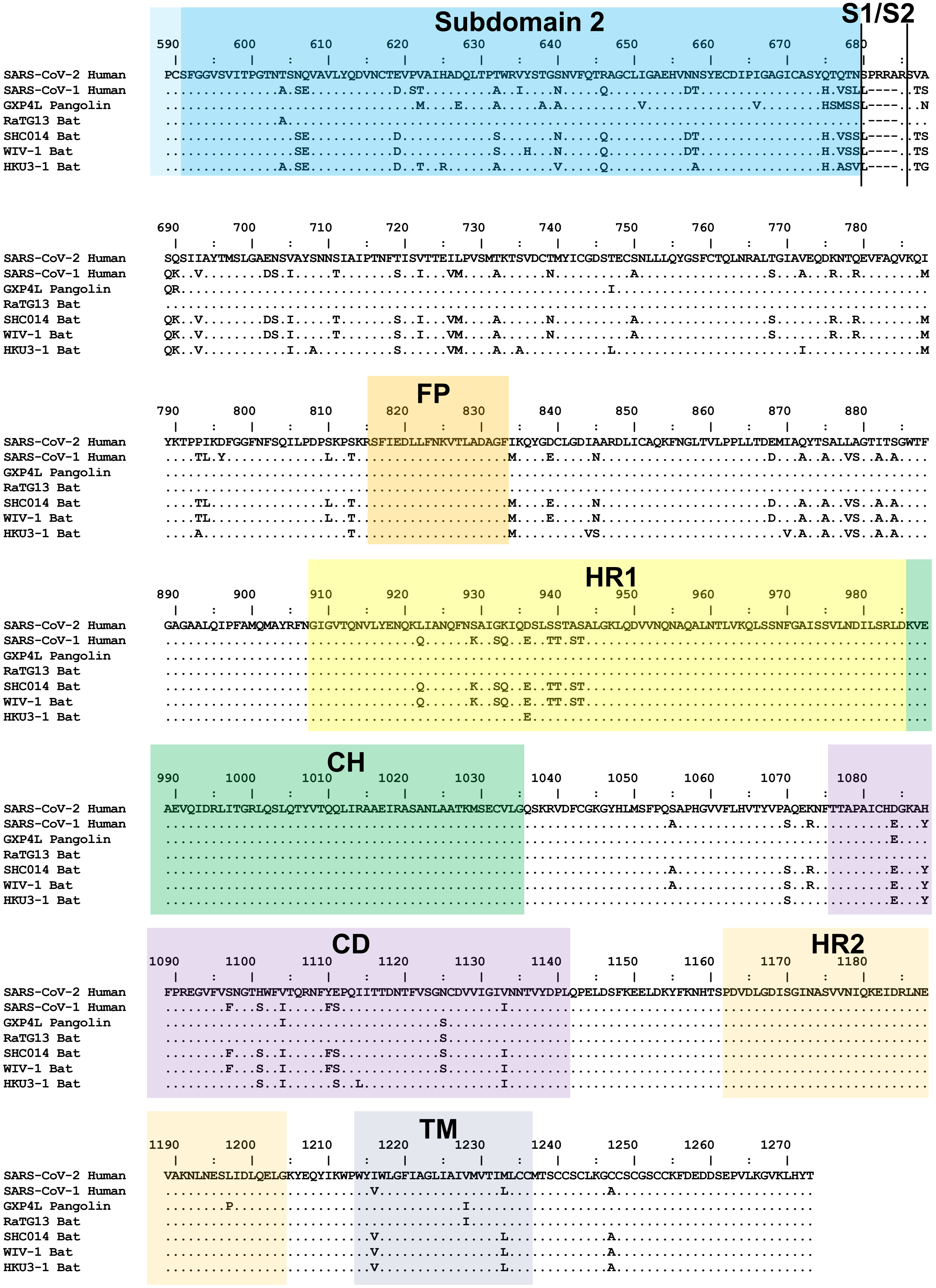

**Extended Data figure 6. Multiple Sequence Alignment of Spike Protein from a Representative Set of Group 2b Betacoronaviruses.**SARS-CoV-2 Wuhan-1 spike protein numbering is shown. NTD=N-terminal domain; RBD=receptor binding domain; S1/S2=SARS2 furin cleavage site; FP=fusion peptide; HR1=heptad repeat 1; HR2=heptad repeat 2; CH=central helix; CD=connecting domain; TM=transmembrane domain.  ACE2 contact positions in SARS2 (calculated from PDB coordinates 6MOJ and 6LZG) are highlighted in dark red.
